# Response Inhibition in Adolescents is Moderated by Brain Connectivity and Social Network Structure

**DOI:** 10.1101/395038

**Authors:** Steven H. Tompson, Emily B. Falk, Matthew Brook O’Donnell, Christopher N. Cascio, Joseph B. Bayer, Jean M. Vettel, Danielle S. Bassett

## Abstract

Self-control is vital for a wide range of outcomes across our lifespan, yet the developmental trajectory of its core components during adolescence remains elusive. Many adolescents can successfully regulate their behavior even when they do not show strong activation in brain regions typically recruited during self-control in adults. Thus, adolescents may rely on other neural and cognitive resources to compensate, including daily experiences navigating and managing complex social relationships that likely bolster self-control processes. Here, we tested whether activity and connectivity in brain systems associated with social cognition (i.e., self-processing and mentalizing) facilitated successful self-control. We measured brain activity using fMRI as 62 adolescents completed a Go/No-Go response inhibition task. Recruitment of social brain systems, especially the self-processing system, was associated with better response inhibition in adolescents. Interestingly, the reliance on the self-processing system was stronger in adolescents with weaker activation in the canonical response inhibition system, suggesting a compensatory role for social brain systems during adolescent development. Furthermore, we examined the importance of social context by computing the size, number of communities, and modularity of our participants’ real-life social network. We found that adolescents with more friends and more communities in their social networks demonstrated a stronger relationship between response inhibition and recruitment of social brain systems. Collectively, our results identify the importance of social context and its moderating role on the relationship between brain activity and behavior. Furthermore, our results indicate a critical role for social brain systems during the developmental trajectory of self-control throughout adolescence.

**Significance Statement:** We employed a network neuroscience approach to investigate the role of social context and social brain systems in facilitating self-control in adolescents. We found that recruitment of social brain systems was associated with better response inhibition in adolescents, especially for adolescents with weaker activation in the response inhibition system. Moreover, adolescents with more friends and communities in their social networks showed stronger relationships between response inhibition and recruitment of social brain systems. Our results advance understanding of how brain systems facilitate self-control in adolescents, and how these brain responses are associated with features of an adolescent’s real-life social network. Bringing together findings related to brain networks and social networks provides key insights into how biology and environment mutually influence development.

## Introduction

Adolescence is characterized by rapid cognitive, social, and neurological development (Blakemore, 2008; Casey, Jones, & Hare, 2008; Ernst, Pine, & Hardin, 2005). In a notable hallmark of the developmental trajectory, the size and structure of the prefrontal cortex changes in concert with an increasing segregation of functional brain networks (Fair et al., 2007). Collectively, these cohesive processes have been posited as the underlying neural substrates to facilitate improved self-control (Casey et al., 2008; Konrad & Eickhoff, 2010; Tamm, Menon, & Reiss, 2002). Response inhibition is a key component of self-control, and deficits in response inhibition, and the brain systems supporting response inhibition, are thought to contribute to altered psychological and behavioral functioning in adolescents, as manifest in ADHD (Konrad & Eickhoff, 2010), substance abuse (Mahmood et al., 2013), and condom use (Hansen, Thayer, Feldstein Ewing, Sabbineni, & Bryan, 2018).

In addition to rapid changes in neurological structure and function, the period of adolescence is also characterized by heightened social sensitivity (Braams & Crone, 2017), and peer influence is a pervasive factor that influences adolescent behavior (Wasylyshyn et al., 2018). Recent work shows that the structure and composition of an adolescent’s social network can influence their self-control (Meldrum, Young, & Weerman, 2012). This putative role of the social network surrounding the individual is particularly notable in light of the fact that self-control can act as an important buffer to risky peer influence (Meldrum, Miller, & Flexon, 2013; Meldrum et al., 2012). In addition, both negative peer influence and reduced tendencies to engage in self-control processes contribute to behavioral issues arising during adolescence (Konrad & Eickhoff, 2010; Meldrum et al., 2013, 2012).

Studies of response inhibition in adults consistently demonstrate that a core set of brain regions are involved in successfully inhibiting prepotent responses, including the right inferior frontal gyrus (IFG), dorsal lateral prefrontal cortex (dlPFC), and basal ganglia (Simmonds, Pekar, & Mostofsky, 2008). Adolescents recruit these brain regions to a lesser extent than adults, and they also show more distributed patterns of activation in other brain regions including the medial prefrontal cortex (mPFC) and posterior cingulate cortex (PCC) (Fair et al., 2007; Marsh et al., 2006; Rubia et al., 2013; Tamm et al., 2002). These studies suggest that recruitment of regions involved in response inhibition facilitates rule-based, top-down control, whereas recruitment of regions in the default mode (DM), including mPFC, PCC, or temporal areas, facilitates bottom-up processing (Konrad & Eickhoff, 2010). Additional evidence suggests that these systems may be less efficient in adolescence than in adulthood, reflecting immature or less developed inhibitory mechanisms (Konrad & Eickhoff, 2010; Marsh et al., 2006; Rubia et al., 2013; Tamm et al., 2002; Vara, Pang, Vidal, Anagnostou, & Taylor, 2014).

Collectively, these results have been interpreted as supporting the notion that recruitment of areas outside of the canonical response inhibition system would lead to deficits in response inhibition (Konrad & Eickhoff, 2010; Tamm et al., 2002). However, it is also possible that activation of a more distributed set of areas reflects an adaptive, compensatory mechanism necessary to support response inhibition in the developing adolescent brain. Adjudicating between these two hypotheses is difficult in part because most fMRI studies on response inhibition in adolescents do not report or do not find significant associations between brain activity and response inhibition performance (Tamm et al., 2002), although some report differential activation in groups of individuals that differ in response inhibition (Liddle et al., 2011).

A network neuroscience framework that also takes into account social context might help address some of these conflicting hypotheses and yield important insights into the neurophysiological drivers of successful response inhibition. One possible explanation for the null result that brain activity is not associated with response inhibition performance is that most studies ignore contextual factors that might moderate the link between brain activity and inhibitory abilities, including social relationships and individuals’ position in their social network (Falk & Bassett, 2017; Pegors, Tompson, O’Donnell, & Falk, 2017; Schmälzle et al., 2017). Recent work suggests that social relationships and an individual’s position in their social network shape how the brain processes information (Schmälzle et al., 2017). Furthermore, social relationships are increasingly being organized into clusters of segregated communities around individuals, such that individuals can now connect with multiple groups that serve different functions (Hampton & Wellman, 2003; Rainie & Wellman, 2012). Two important features of this individual-focused organization of social networks are that (i) it requires individuals to actively recruit other individuals into their network and maintain existing communities within their network, and (ii) it allows individuals to keep groups more segregated for distinct functions (e.g., one group provides advice about social and romantic relationships, another group provides support for school and career issues; Rainie & Wellman, 2012). Thus, one might expect that the degree to which an individual’s social network is organized into many (versus few) or segregated (versus overlapping) communities would be associated with that individual’s cognitive abilities, including what cognitive or social resources individuals are likely to recruit to regulate their behavior. We suggest that adolescents with larger networks segregated into more communities may adopt more social resources to successfully regulate their behavior.

Another possible explanation for the null result that brain activity is not associated with response inhibition performance is that the cognitive demands of a response inhibition task may require the coordinated action of multiple brain regions and systems (Chai et al., 2017; Shine et al., 2016), making connectivity phenotypes critical for predicting response times (Vatansever, Menon, Manktelow, Sahakian, & Stamatakis, 2015). Although some evidence suggests that behavioral issues in adolescents may arise from weaker activation in control systems compared to adults, as well as more distributed activation elsewhere in the brain (Konrad & Eickhoff, 2010), it is also possible that adolescents compensate for weaker recruitment of control systems by leveraging social resources or recruiting other cognitive control systems. Given adolescents’ sensitivity to social influence (Braams & Crone, 2017; Wasylyshyn et al., 2018), we suggest that social brain systems might be particularly important for understanding how adolescents recruit self-control processes.

Here, 62 adolescent males completed a Go/No-Go response inhibition task while brain activity was measured using fMRI. We also collected information about adolescents’ real-life social networks in order to assess the moderating role of social network properties in influencing the link between brain activity and response inhibition. We hypothesize that distributed patterns of activation across both response inhibition brain regions as well as other socially-relevant brain systems (self-processing and mentalizing systems) should be associated with response inhibition performance, and this distributed neural activity will be moderated by social network properties. This hypothesis is based on the notion that for some individuals, recruiting regions outside the canonical response inhibition network might also be important for effective response inhibition. Specifically, an individual’s social network might play an important role in influencing the cognitive strategies and networks recruited to successfully complete the Go/No-Go task. We anticipate that social brain systems, including brain systems involved in self-referential processing and mentalizing, should facilitate response inhibition, especially for adolescents with weaker recruitment of executive function systems. Additionally, we expect that these effects will be moderated by an adolescent’s real-life social network structure, including network size, number of communities, and network modularity.

## Results

### Behavioral Performance on a Task Requiring Response Inhibition

We first examined participants’ behavioral performance on the task requiring response inhibition. Participants’ average response time on Go trials was 373 ms (*SD*=4.36 ms), and their average accuracy on No-Go trials was 75.4% (*SD*=11.0%). Consistent with past work, we observed a speed-accuracy tradeoff such that participants who showed greater accuracy on No-Go trials responded more slowly on Go trials (*r*(60)=0.373, *p*=0.003). For all subsequent analyses, we focused on a metric known as Go/No-Go efficiency, which quantifies how well participants balanced the speed-accuracy tradeoff when completing the Go/No-Go task. The Go/No-Go efficiency score ranges from 0-1, where values approaching 1 indicate that participants are responding quickly but still correctly inhibiting responses on No-Go trials, and values approaching zero indicate that a participant was either responding fast but inaccurately, or slow and accurately. Participants’ average task efficiency was 0.497 (*SD*=0.081).

### Activation of Brain Systems During a Task Requiring Response Inhibition

We next examined recruitment of the three brain systems of interest (the response-inhibition, self-processing, and mentalizing systems) during trials that required response inhibition. As shown in panel B of Figure 1, mean system activation was significantly greater during correct No-Go trials than during correct Go trials in both the response inhibition system (*t*(60)=3.58, *p*<0.001) and the mentalizing system (*t*(60)=5.51, *p*<0.001), but brain activity was not greater on average during response inhibition in the self-processing system (*t*(60)=1.25, *p*=0.216).

**Figure 1.**
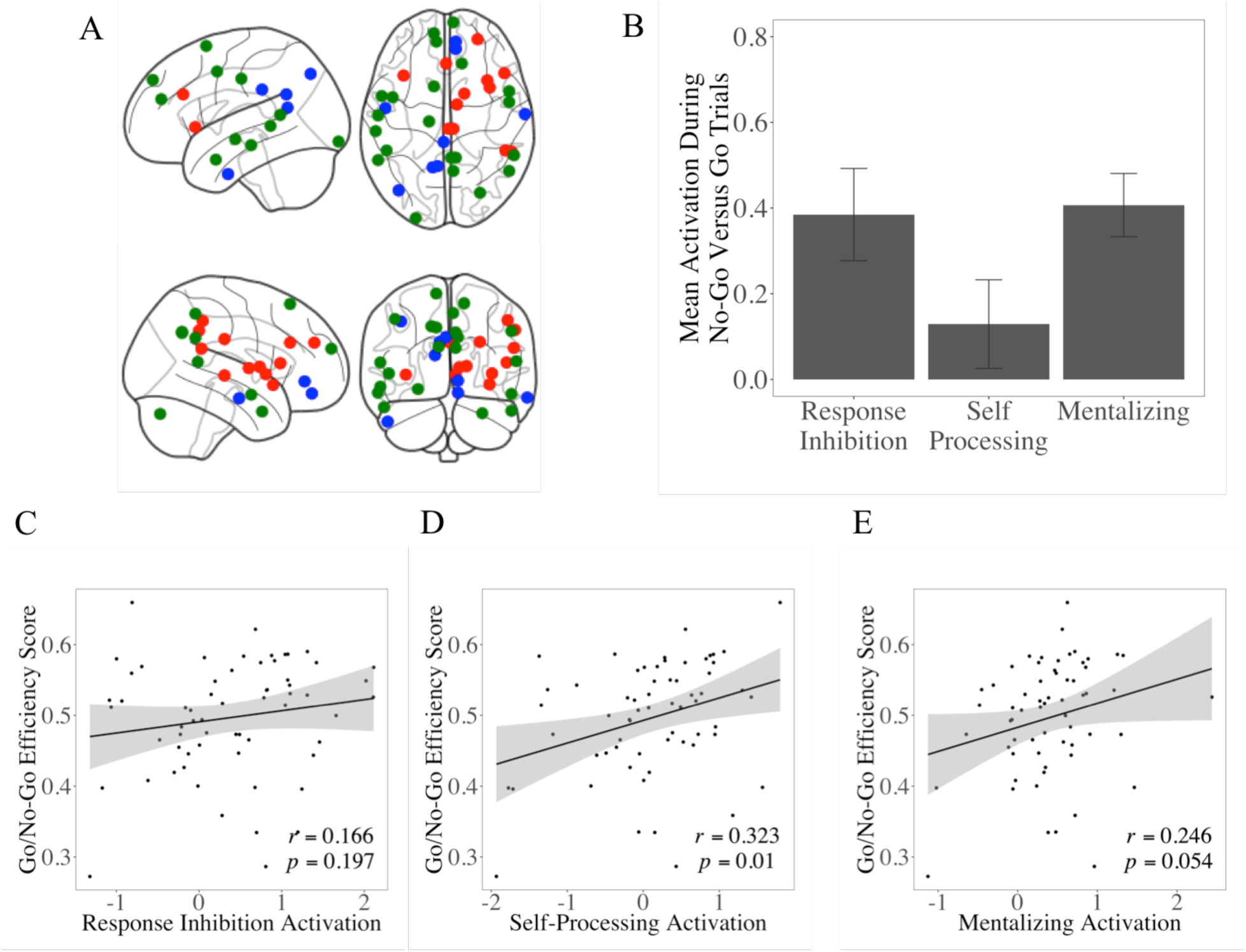
Recruitment of brain systems during a task requiring response inhibition. (*A*) Regions in the canonical response inhibition system identified using Neurosynth included the basal ganglia and IFG (red), whereas self-processing regions included the ventral mPFC, PCC, and temporal pole (blue), and mentalizing regions included the dorsal mPFC, PCC, and TPJ (green). (*B*) Average activation in the response inhibition, self-processing, and mentalizing systems for correct No-Go trials versus correct Go trials. (*C*) Scatterplot of Go/No-Go efficiency score versus mean activation of the response inhibition system. (*D*) Scatterplot of Go/No-Go efficiency score versus mean activation of the self-processing system. (*E*) Scatterplot of Go/No-Go efficiency score versus mean activation of the mentalizing system.

To test how brain activity related to the tradeoff between speed and accuracy when inhibiting prepotent responses in this task, we examined whether mean activation for No-Go trials versus Go trials in each of the three systems accounted for variability in the adolescents’ Go/No-Go efficiency score (see Methods, Eq. 1). Interestingly, brain activity in the response inhibition system was not correlated with individual differences in efficiency (*r*(60)=0.166, *p*=0.197; Figure 1 panel C), but greater recruitment of the self-processing system was positively associated with efficiency (*r*(60)=0.323, *p*=0.010; Figure 1 panel D). The mentalizing system showed a similar trend as the self-processing system, but its activation was also marginally associated with better efficiency (*r*(60)=0.246, *p*=0.054; Figure 1 panel E).

### Compensatory Activation in Social Brain Systems

We next examined our hypothesis that adolescents with less tendency to use executive function brain systems typically observed in adults may instead recruit regions outside the canonical response inhibition network to successfully perform the task. We ran two multiple regression analyses between the response inhibition network and each of the two social brain systems, with mean system activation for No-Go versus Go trials as the independent variables and the Go/No-Go efficiency score as the dependent variable.

We found a significant interaction between response inhibition activation and self-processing activation: adolescents who had weaker activation of the response inhibition system showed a stronger relationship between task performance and activation of the self-processing system (*β*=-0.214, *p*=0.035; Figure 2 panel A). To further explore this interaction effect, we probed the simple slopes: at lower levels of response inhibition system activation (−1 SD), the relationship between self-processing system activation and efficiency was significant (*β*=0.487, *p*=0.001). At higher levels of response inhibition system activation (+1 SD), there was no significant relationship between self-processing system activation and efficiency (*β*=0.059, *p*=0.722).

**Figure 2.**
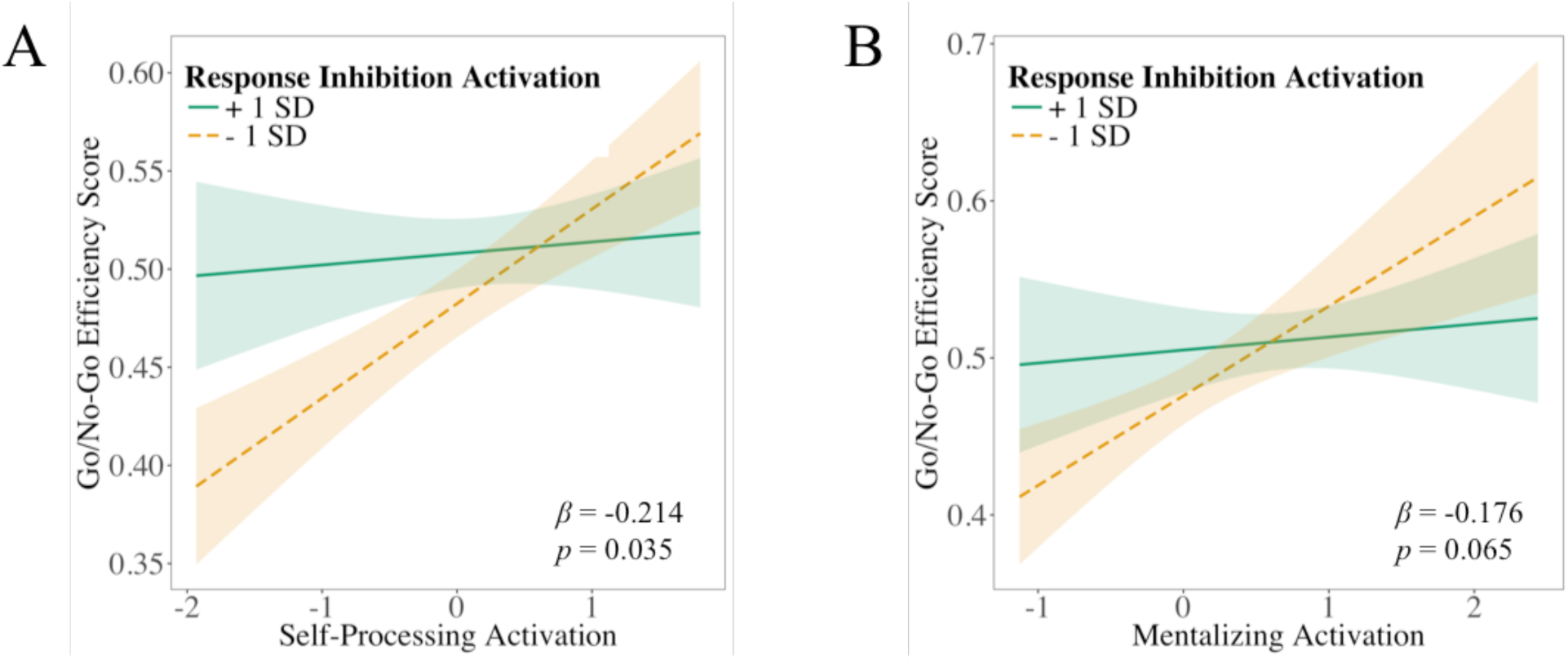
Interaction between response inhibition activation and social brain systems. (*A*) Relationship between mean activation in the self-processing system and Go/No-Go efficiency scores differs for adolescents with stronger mean activation in the response inhibition system (solid line) versus adolescents with weaker mean activation in the response inhibition system (dashed line). (*B*) Relationship between mean activation in the mentalizing system and Go/No-Go efficiency scores differs for adolescents with stronger mean activation in the response inhibition system (solid line) versus adolescents with weaker mean activation in the response inhibition system (dashed line).

We found a similar, albeit marginal, effect with response inhibition system activation and mentalizing system activation: adolescents who had weaker activation of the response inhibition system showed a stronger relationship between task performance and activation of the mentalizing system (*β*=-0.176, *p*=0.065; Figure 2 panel B). However, after examining simple slopes, we found a compensatory relationship at lower levels of response inhibition system activation. The relationship between mentalizing system activation and efficiency was significant at lower levels (−1 SD) of response inhibition system activation (*β*=0.412, *p*=0.022), but not at higher levels (+1 SD) of response inhibition system activation (*β*=0.060, *p*=0.717).

Across both regression analyses, activation in self-processing and mentalizing regions facilitated a higher Go/No-Go efficiency score only when adolescents had lower activation in response inhibition regions, suggesting that these regions might serve a compensatory role for adolescents with less mature brain development.

### Connectivity Within and Between Brain Systems and Response Inhibition

In addition to examining mean system activation, we were also interested in determining whether connectivity within and between these three systems was associated with successfully and efficiently inhibiting prepotent responses. If self-processing and mentalizing systems are compensating for weaker recruitment of response inhibition systems, then it is possible that communication between these regions is important for efficient response inhibition. To assess the role of connectivity in this process, we examined the average connectivity both *within* each of the three systems as well as the average connectivity *between* the response inhibition network and each of the social brain systems (self-processing and mentalizing). For both within-and between-system comparisons, we computed the Pearson correlation coefficient between average connectivity and task performance (see Figure 3).

**Figure 3.**
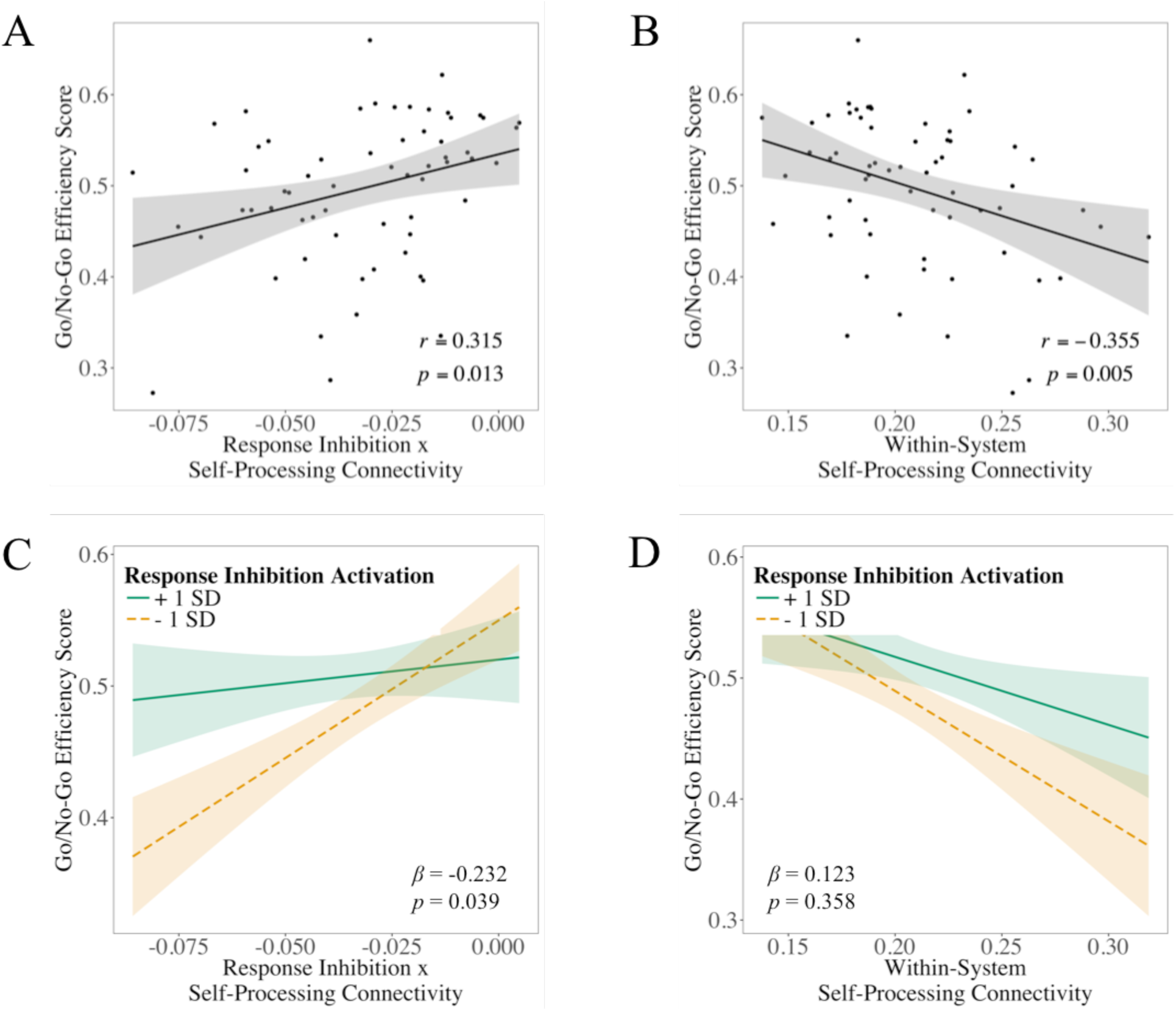
Relation between inter-system connectivity and Go/No-Go efficiency score. (*A*) Scatterplot of Go/No-Go efficiency score versus connectivity between the response inhibition system and self-processing system. (*B*) Scatterplot of Go/No-Go efficiency score versus connectivity within the self-processing system. (*C*) Relationship between response inhibition system x self-processing system connectivity and Go/No-Go efficiency scores differs for adolescents with stronger activation in the response inhibition system (solid line) versus adolescents with weaker activation in the response inhibition system (dashed line). (*D*) Relationship between connectivity within the self-processing system connectivity and Go/No-Go efficiency scores does not differ for adolescents with stronger activation in the response inhibition system (solid line) versus adolescents with weaker activation in the response inhibition system (dashed line).

We found that adolescents who had a higher Go/No-Go efficiency score had stronger connectivity between the response inhibition and self-processing systems (*r*(60)=0.315, *p*=0.010; Figure 3 panel A) and weaker connectivity within the self-processing system (*r*(60)=-0.355, *p*=0.005; Figure 3 panel B). We observed no significant associations between mentalizing system connectivity and task performance.

### Unique Contribution of Mean Activity and Average Connectivity for Response Inhibition

Since both the mean system activation and connectivity results demonstrated a compensatory role for the social brain systems, we included both measurements of brain activity in the same model to understand their relationship to improved performance on the response inhibition task.

We found a significant interaction between response inhibition activation and the connectivity between the response inhibition system and the self-processing system (*β*=-0.232, *p*=0.039; Figure 3 panel C). To further explore this interaction effect, we probed the simple slopes and found that at lower levels of response inhibition system activation (−1 SD), the relationship between response inhibition × self-processing system connectivity and an adolescent’s Go/No-Go efficiency score was significant (*β*=0.559, *p*=0.001). At higher levels of response inhibition system activation (+1 SD), there was no significant relationship between response inhibition × self-processing system connectivity and the efficiency score (*β*=0.096, *p*=0.553).

Interestingly, we observed no such moderating effect for connectivity within the self-processing system (*β*=0.123, *p*=0.358; Figure 3 panel D). Collectively, these results suggest that activation in the self-processing system supports efficient task performance when it involves communication with response inhibition brain regions, but not when it involves communication with other self-processing brain regions.

### Real-life Social Network Properties Correlate with the Recruitment of Brain Systems

Across our analyses, results demonstrated that activation in the two social brain systems (self-processing and mentalizing) compensated for weaker recruitment of the canonical response inhibition system, and this compensatory activity was associated with higher Go/No-Go efficiency scores, enabling enhanced task performance. In short, brain systems implicated in social processes facilitate better response inhibition performance. Consequently, our next set of analyses examined whether features of an adolescent’s real-life social network correlated with the neural activation when adolescents successfully inhibited prepotent responses.

We computed three features of the adolescents’ social networks: the number of communities, the size of communities (number of friends in a community), and the modularity of the network that characterize the interrelations among the communities. A network with high modularity contains distinct communities of friends where the connections within communities are substantially denser than the connections between communities. In contrast, a network with low modularity may also contain distinct communities, but with less of a difference in the density of connections within versus between communities. An adolescent with high network modularity may have sets of friends from school, another from sports, and yet another from church, and the friends in those groups may never interact with one another. An adolescent with low network modularity might have friends from these same groups, but friends from school might also know friends from sports or church. In our study, adolescents had an average of 491 Facebook friends in their social network (*SD*=280), which were clustered into an average of 8.42 communities (*SD*=4.07) with a mean modularity of 0.235 (*SD*=0.122).

For each of these three social network parameters, we investigated whether variability in the adolescent’s neural response across the three brain systems during the response inhibition task successfully accounted for variability observed in their real-life social network. This set of analyses separately explored the relationship of the social network parameters with each of the brain parameters identified from the previous results: mean system activation of each system, within-system connectivity of each system, and between-system connectivity of the response inhibition system and each of the two social brain systems.

We found that adolescents with more friends in their social network had greater connectivity within the mentalizing system (*r*(60)=0.259, *p*=0.042; Figure 4 panel A). A similar, but only marginally significant, relationship was observed with mean system activation, where adolescents who had more communities in their social network also had greater activation within the mentalizing system (*r*(60)=0.224, p=0.080; Figure 4 panel B). Finally, we observed a negative correlation between within-system response inhibition connectivity and network modularity; adolescents with weaker connectivity within the response inhibition system had more segregated communities in their social network than adolescents with stronger connectivity within the response inhibition system (*r*=-0.351, *p*=0.005). None of the other correlations were significant.

**Figure 4.**
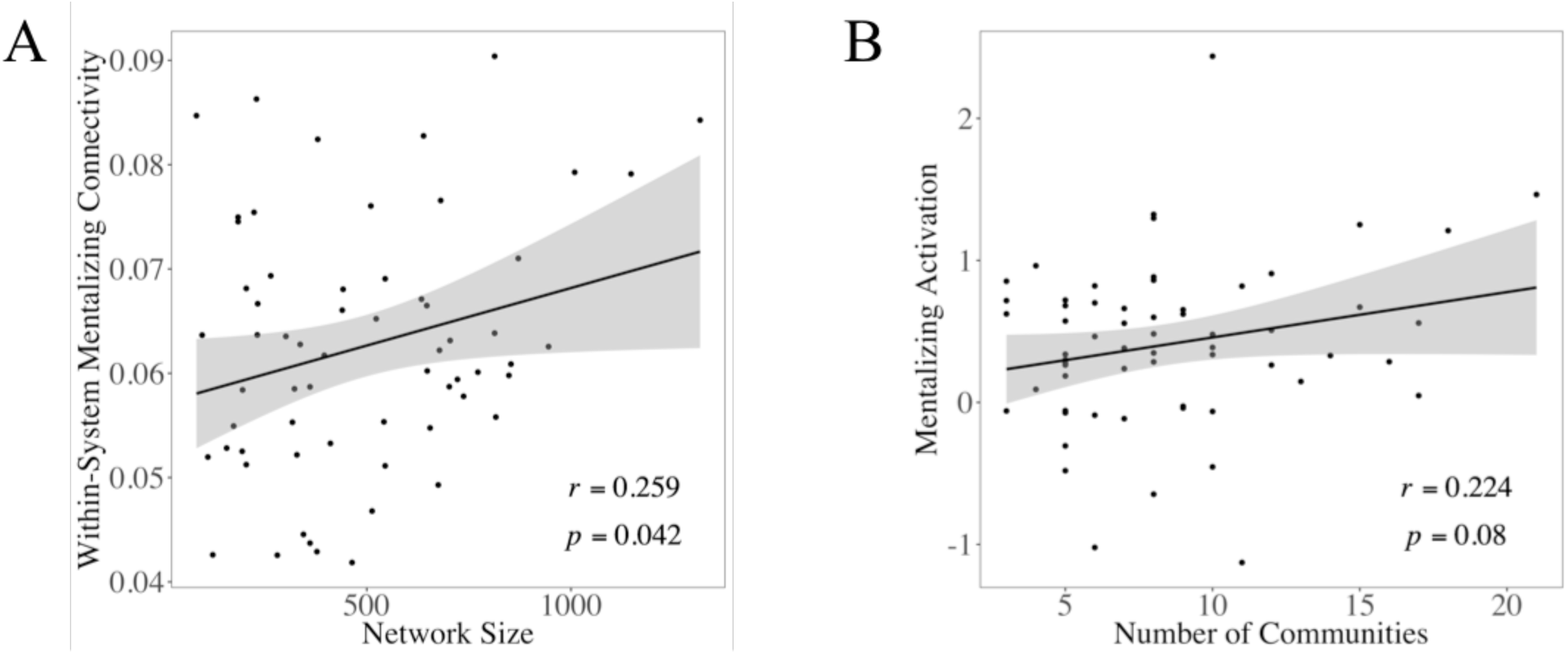
Association between mentalizing system and social network properties. (*A*) Scatterplot of social network size (number of friends) and connectivity within the mentalizing system. (*B*) Scatterplot of number of social network communities and mean activation in the mentalizing system.

### Real-life Social Networks Account for Compensatory Role of Social Brain Systems

In our final set of analyses, we examined whether any of the three social network properties (number, size, and modularity of communities) moderated the observed relationship between brain activity or connectivity and adolescents’ Go/No-Go efficiency score. Mirroring our correlational analyses, we tested each of these hypotheses in separate models for the mean activity, within-system connectivity, and between-system connectivity of the three brain systems of theoretical interest: response inhibition, mentalizing, and self-processing systems.

We found that the relationship between mean activity in the response inhibition system and an adolescent’s Go/No-Go efficiency score was moderated by social network properties. Specifically, there was a significant interaction between mean activity in the response inhibition system and network modularity (*β*=0.344, *p*=0.026). Additionally, there were two marginally significant interactions: one occurred between activity in the response inhibition system and the number of communities (*β*=0.299, *p*=0.085; Figure 5 panel A), whereas the other occurred between activity in the response inhibition system and the network size (*β*=0.211, *p*=0.084). We further probed these interactions using simple slopes analysis. We found that adolescents high (+1 SD) in network size, number of communities, and modularity had a significant positive relationship between mean activity in the response inhibition system and the Go/No-Go efficiency score (*β*=0.390, *p*=0.027; *β*=0.530, *p*=0.034; *β*=0.625, *p*=0.009; respectively), whereas the relationship was not significant for adolescents low (−1 SD) in network size, number of communities, and modularity (*β*=-0.033, *p*=0.853; *β*=-0.068, *p*=0.713; *β*=-0.062, *p*=0.703; respectively). Interestingly, no significant interactions were found between these three social network properties and mean activation in either of the two social brain systems.

**Figure 5.**
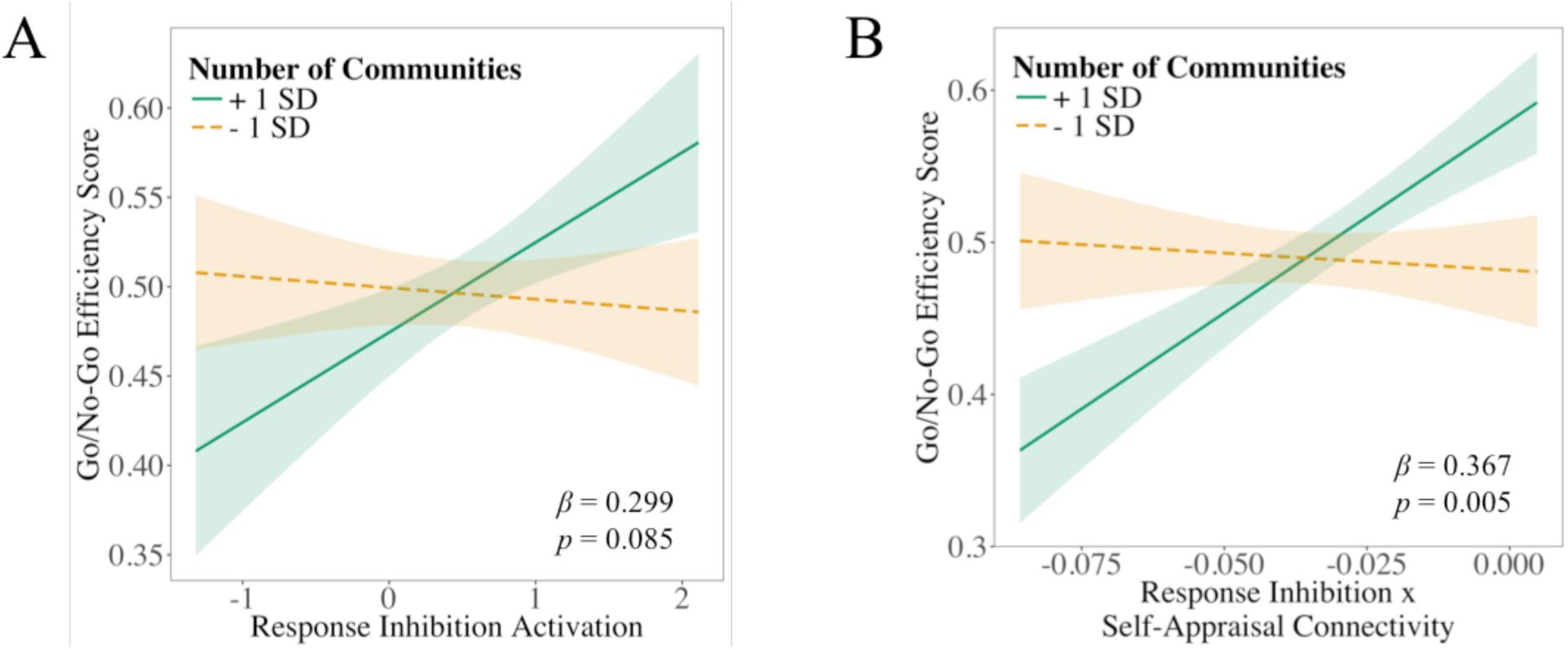
Social network properties moderate the relationship between brain and behavior. (*A*) Relationship between response inhibition system x self-processing system connectivity and Go/No-Go efficiency scores differs for adolescents with more communities in their social network (solid line) versus adolescents with fewer communities in their social network (dashed line). (*B*) Relationship between activation in the response inhibition system and Go/No-Go efficiency scores differs for adolescents with more communities in their social network (solid line) versus adolescents with fewer communities in their social network (dashed line).

In contrast, the connectivity results revealed that an adolescent’s real-life social network accounted for the compensatory role that social brain systems served for better task performance. More specifically, the association between brain connectivity and task performance was significantly moderated by the number of social network communities, but not by social network size or modularity. The number of communities significantly moderated the relationship between the Go/No-Go efficiency score and the between-system connectivity of the response inhibition and self-processing systems (*β*=0.367, *p*=0.005; Figure 5 panel B). Compared to adolescents with fewer communities in their social networks (−1 SD; *β*=-0.068, *p*=0.713), adolescents with more communities (+1 SD) showed a stronger positive association between their Go/No-Go efficiency score and the between-system connectivity of the response inhibition and self-processing systems (*β*=0.530, *p*=0.034).

We also observed a marginally significant moderating effect of number of communities on the relationship between the Go/No-Go efficiency score and within-system mentalizing connectivity (*β*=-0.250, *p*=0.058). Adolescents with a large number of communities exhibited a negative relationship between task performance and within-system mentalizing connectivity, whereas adolescents with few communities showed a positive relationship between their Go/No-Go efficiency score and within-system mentalizing connectivity; however, the simple slopes were not significant in either case (*β*=-0.277, *p*=0.134; *β*=0.224, *p*=0.224; respectively).

## Discussion

Self-control processes, including response inhibition, predict many important outcomes in adolescence, including school success (Blair & Diamond, 2008), risky behaviors (Behan et al., 2014; Hansen et al., 2018; Mahmood et al., 2013), and psychiatric outcomes (Konrad & Eickhoff, 2010; Liddle et al., 2011). Social context also influences how adolescents engage self-control processes, such that family and peer relationships can facilitate better self-control or buffer against potential negative effects of weaker self-control (Farley & Kim-Spoon, 2014; Meldrum et al., 2012). Yet, the neurophysiological drivers of successful response inhibition in adolescents remain unclear. We argue that progress in understanding has been hampered in part by a focus on activation in single brain regions as well as a lack of focus on the social context surrounding the adolescent.

Here, we adopted a network neuroscience approach (Bassett & Sporns, 2017; Bullmore & Sporns, 2009) to examine how response inhibition, self-processing, and mentalizing systems contribute to effective response inhibition. We also collected adolescents’ real-life social networks to examine whether variability in social network structure could account for differences in adolescent’s ability to inhibit prepotent responses. We found that brain regions outside the canonical response inhibition system compensated for adolescent differences in the response inhibition system, such that adolescents who had weaker response inhibition activity still performed well on the Go/No-Go task if they had stronger activity in social brain regions and greater connectivity between social brain regions and the response inhibition network during the task. Moreover, adolescents with larger social networks with segregated communities of friends showed stronger relationships between brain systems and response inhibition. Collectively, our results provide insight into how brain systems facilitate cognitive control in adolescents, and how these brain responses are associated with features of an adolescent’s real-life social network.

### Activation in Social Brain Systems

This study extends previous work that finds more distributed patterns of brain activity in adolescents during response inhibition (Fair et al., 2007; Marsh et al., 2006; Rubia et al., 2013; Tamm et al., 2002). We showed that aggregate activity in the canonical response inhibition system as well as aggregate activity in the mentalizing system was significantly greater during No-Go trials than during Go trials, yet performance on the task, as indexed by the Go/No-Go efficiency score, was significantly associated with greater activation in the self-processing system. Although we found no direct relationship between activation in the response inhibition system and how efficiently adolescents inhibited their responses on No-Go trials, we found that this relationship was moderated by network dynamics in social brain systems including self-processing and mentalizing systems. We showed here for the first time that adolescents who have weaker response inhibition activation still perform well on the Go/No-Go task if they have stronger activation in self-processing and mentalizing systems, and if they have greater connectivity between the self-processing and response inhibition system during the task.

The pattern of results that we uncovered is consistent with past work that has also found that adolescents show weaker activation in response inhibition brain regions (e.g., basal ganglia and IFG) and stronger activation in social brain regions (e.g., mPFC and PCC; Fair et al., 2007; Marsh et al., 2006; Rubia et al., 2013; Tamm et al., 2002). These prior studies have also found weak evidence for a relationship between activation in response inhibition brain regions and task performance (Tamm et al., 2002). The current work suggests that one reason for this lack of a direct effect is that other more distributed brain systems might be compensating for weaker recruitment of executive function in the adolescent brain.

Importantly, the version of the Go/No-Go task used in the current study has no explicit social components, and so one might wonder how or why social brain systems are recruited during a non-social task. Since most brain regions serve multiple functions, one possibility is that the cortical regions implicated in social processing are being co-opted for response inhibition to compensate for weaker activation in other brain regions. It is also possible that, because adolescents are highly sensitive to social information, they are recruiting additional motivational or cognitive strategies from their daily social experiences to facilitate effective task performance (e.g., to perform well in front of the experimenter, feel proud of their performance, or implicitly compete against others in the study). Another possibility is that self-control processes may first develop specifically for social situations in social brain systems, and then self-control becomes more domain-general once the canonical response inhibition system observed in adults gradually develops over the course of adolescence.

### Connectivity between Response Inhibition and Social Brain Systems

If self-processing and mentalizing systems are compensating for weaker recruitment of response inhibition systems, then it is possible that communication between these regions is important for efficient response inhibition. Consistent with this idea, we found that connectivity between the response inhibition and self-processing systems was positively associated with response inhibition performance. Importantly, connectivity within the self-processing system was negatively associated with response inhibition performance, suggesting that greater recruitment of social brain systems is not always beneficial for response inhibition. Thus, social brain systems may help compensate for weaker recruitment of response inhibition systems, but only when social brain systems are communicating directly with more canonical response inhibition brain regions.

Recent work in network neuroscience also suggests that successful performance on many cognitive tasks requires coordinated action across multiple brain regions and brain systems (Chai et al., 2017; Shine et al., 2016). This role for distributed connectivity extends to regions that are not typically considered important for a specific cognitive process, many of which overlap with the regions identified in our self-processing and mentalizing meta-analyses. For example, regions in the default mode network facilitate faster response times in a motor task (Vatansever et al., 2015) and better working memory performance (Čeko et al., 2015), but are not canonically thought of as part of motor or memory circuitry. Similarly, we show here that brain systems implicated in social processes facilitate better response inhibition performance. Notably, our study is the first to show this relationship in adolescents.

### Social Network Structure and Response Inhibition

The current work also contributes to a growing body of evidence that social network structure influences neural processes (Bickart, Hollenbeck, Barrett, & Dickerson, 2012; Molesworth, Sheu, Cohen, Gianaros, & Verstynen, 2015; O’Donnell, Bayer, Cascio, & Falk, 2017; Pegors et al., 2017; Powell, Lewis, Roberts, Garcia-Finana, & Dunbar, 2012; Schmälzle et al., 2017). The structure of and activation in brain regions implicated in social processing and affective processing are influenced by individuals’ social network size (Kanai, Bahrami, Roylance, & Rees, 2012; Lewis, Rezaie, Brown, Roberts, & Dunbar, 2011; Powell et al., 2012; Von Der Heide, Vyas, & Olson, 2014). Social network structure also influences brain activity in mentalizing regions when thinking about others’ opinions (O’Donnell et al., 2017) as well as connectivity within the mentalizing system during a social exclusion task (Schmälzle et al., 2017).

Here, we showed for the first time that social network structure also moderated brain systems involved in a non-social task. Adolescents with larger social networks comprised of a greater number of segregated communities showed a stronger relationship between response inhibition activation and task performance. Adolescents with more communities in their social network also had a stronger relationship between task performance and the between-system connectivity of the response inhibition and self-processing systems. Larger social network structures that have more communities or communities that are more segregated (higher in modularity) require individuals to actively maintain multiple groups of friends (Hampton & Wellman, 2003; Rainie & Wellman, 2012). The ability to actively maintain these groups may in turn be facilitated by (or require) more diverse brain systems being recruited for behavioral self-regulation.

This pattern of results is consistent with recent work showing that peer relationships and social context can strongly influence self-control. Children’s ability and motivation to regulate their behavior is influenced by salient group norms (Doebel & Munakata, 2018), and adolescents who are surrounded by peers with better self-reported self-control are also more likely to show improved self-control over the course of adolescence (Meldrum et al., 2012). Brain activity during a social exclusion task in the laboratory also predicts an adolescent’s susceptibility to peer influence in a driving simulation one week later (Wasylyshyn et al., 2018). Here, we showed that an adolescent’s social network structure also influences which brain systems serve to help regulate their behavior. It is possible that adolescents’ daily experiences navigating and managing complex social relationships with multiple distinct communities influence how they use different cognitive strategies or motivational resources to complete cognitive tasks such as response inhibition. Alternatively, adolescents who recruit more diverse brain systems may be more capable of managing and maintaining larger and more complex social networks. An interesting avenue for future work would be to examine the directionality of this relationship between social brain network processing and the structure of real-life social networks.

### Methodological Considerations and Limitations

One potential limitation for the generalizability of the current work concerns the sample. All of our participants were 16 year-old males, and thus it is unclear whether these results would extend to adolescent females as well. The original study for which these data were collected was primarily concerned with neural correlates of adolescent risky driving and was restricted to adolescent males who had recently received their driver’s license since this group has the highest statistical risk for accidents on modern roadways.

Since participation was restricted to 16 year-olds, we also cannot address any questions related to the developmental trajectory of response inhibition over the course of adolescence. The adolescent brain is rapidly developing over time, as is adolescents’ response inhibition performance, and thus, future work could investigate whether self-processing and mentalizing systems also compensate for weaker recruitment of response inhibition systems when younger adolescents engage in response inhibition. Previous work has also found that the developmental trajectories of brain activity during response inhibition differ for young males and females (Rubia et al., 2013) and therefore it is also possible that there might be an interaction between age and gender that influences the neural mechanisms underlying response inhibition in adolescents. It would be interesting in future work to investigate this possibility.

Finally, our social network metrics were constructed based on adolescents’ Facebook relationships, and this is only one method to identify social relationships. There are likely other types of relationships that influence adolescents’ self-control that are poorly captured by online social network metrics, such as relationships with teachers or other adult family members who are not linked through online social media. Future work using multiple approaches to collect information about social networks (c.f., Vettel et al., 2018) might yield further insights into the link between brain networks and social networks and their importance for adolescent development.

## Conclusions

In the current work, we employed a network approach to analyze brain data and examined the moderating role of social context. Taken together, this work suggests that adolescents with larger social networks with more communities recruit more diverse brain systems to successfully inhibit prepotent responses. Our results demonstrate that the relationship between behavior and brain activity, as well as connectivity between brain systems, is context-dependent. These results strongly motivate future work to examine how elements of the social context influence adolescent brains. Our work also provides insight into developmental differences in neural mechanisms involved in response inhibition. Previous work has found that adolescents, relative to adults, are more likely to recruit brain regions outside the canonical response inhibition network when completing a Go/No-Go task (Rubia et al., 2013; Tamm et al., 2002). Although this observation is often explained by an assumption of less mature or less developed brain states reflective of weaker executive function in adolescents, we suggest that these more distributed brain activations might instead reflect an adaptive response to an adolescent’s social environment.

## Method

### Participants

One hundred and three adolescent males who recently received their license (restricted to 16-year-olds) were recruited through the Michigan state driver registry database as part of a larger study on peer influences on adolescent driving. Participants met standard MRI safety criteria. In accordance with Institutional Review Board approval from the University of Michigan, legal guardians provided written informed consent, and adolescents provided written assent. Due to technical malfunction, social network data was lost for 29 participants and Go/No-Go performance data was lost for an additional 10 participants. One participant was excluded due to poor performance on the Go/No-Go task (accuracy on No-Go trials less than 50%), and one participant was excluded due to excessive head motion during the fMRI scan (greater than 3mm framewise displacement). Consequently, analyses were conducted on the 62 remaining 16-year-old adolescent males.

### Response Inhibition Task (Go/No-Go)

While lying prone in a scanner continuously acquiring BOLD MRI data, adolescents completed a series of tasks including a response inhibition Go/No-Go task (Aron, Fletcher, Bullmore, Sahakian, & Robbins, 2003; Logan, 1994). In this task, 80% of trials were considered Go trials and 20% of trials were considered No-Go trials. On each trial, a letter was presented on the screen. Participants were instructed to press a button if the letter was an A through F, but they had to withhold their response and not press a button if the letter was an X. Letters were presented for 500 ms, followed by a 1000 ms fixation interval. Go trials were considered correct if participants pressed a button before the next trial began, whereas No-Go trials were considered correct if participants did not press a button before the next trial began. To account for the tradeoff between speed and accuracy when inhibiting prepotent responses, performance was measured using Go/No-Go efficiency:

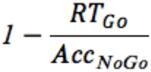

Where *RT*_*Go*_ represents the average response time on Go trials and *Acc*_*NoGo*_ represents the percentage of No-Go trials where the participants correctly withheld a response. To enhance interpretability, we subtracted the proportion of response time to accuracy from unity so that higher scores indicate better performance.

### Real-life Social Network Properties

Outside of the scanner, participants provided access to their Facebook network data using an online survey. Specifically, participants logged into their Facebook account, and the Facebook OpenGraph API (collected in 2011-2013) was used to assess participants’ Facebook activity, friends, and links between friends. We first anonymized the data and then used the NetworkX package implemented in Python 2.7 to construct binary, undirected graphs of each participant’s social network where each Facebook friend is represented as a node in the graph and each connection between friends (Facebook friendship) is represented as an edge on the graph. Each graph was encoded in an N × N adjacency matrix where N is the number of Facebook friends and the ij^th^ matrix element represents whether person *i* is a friend of person *j*.

Using NetworkX, we then computed the size, number, and modularity of communities for each adolescent’s social network. The size of each network was defined as the total number of friends. To determine the number of communities in each network, we used a Louvain-like locally greedy algorithm (Blondel, Guillaume, Lambiotte, & Lefebvre, 2008) to maximize the following modularity quality function:

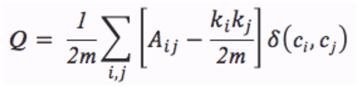

where *A*_*ij*_ represents the strength of the connection between nodes *i* and *j, k*_*i*_ represents the sum of the connection strengths for nodes connected to node *i, 2m* is the sum of all the connection strengths in the network, δ is the Kronecker delta, and *c*_*i*_ represents the community to which node *i* is assigned. Intuitively, the algorithm segregates friends into non-overlapping groups by maximizing the number of within-group connections, in comparison to that expected in an appropriate random network null model, which here we define as the configuration model (also known as the Newman-Girvan null model; Newman & Girvan, 2004).

After optimizing the modularity quality function, we obtain both a partition of nodes into communities and a maximum Q value, the latter of which is often referred to as the network modularity (Newman, 2006). Network modularity ranges from 0 to 1, where a densely connected network (i.e., all of an adolescent’s friends are also friends with each other) has a score closer to 0 whereas a segregated network (i.e., an adolescent has separate clusters of friends from school, the neighborhood, sports) has a score closer to 1.

### fMRI Data Acquisition and Preprocessing

Functional images were recorded using a reverse spiral sequence (repetition time = 2,000 ms, echo time = 30 ms, flip angle = 90°, 43 axial slices, field of view = 220 mm, slice thickness = 3 mm, voxel size = 3.44 × 3.44 × 3.0 mm). We also acquired in-plane T1-weighted images (43 slices, slice thickness = 3 mm, voxel size = 0.86 × 0.86 × 3.0 mm) and high-resolution T1-weighted images [spoiled gradient recall (SPGR) acquisition, 124 slices, slice thickness = 1.02 × 1.02 × 1.2 mm] for use in co-registration and normalization. Functional data were preprocessed using Statistical Parametric Mapping (SPM8, Wellcome Department of Cognitive Neurology, Institute of Neurology, London). The first four volumes were discarded before analysis. Functional images were despiked using the 3dDespike program as implemented in the AFNI toolbox, corrected for differences in slice time acquisition, and spatially realigned to the first functional image. To mitigate remaining nuisance signals, we then applied a high-pass filter with a cutoff of 128 sec, and the subsequent volumes were weighted according to the inverse of their noise variance using the robust weighted least squares toolbox (Diedrichsen, Hashambhoy, Rane, & Shadmehr, 2005). Functional and structural images were co-registered using a two-stage procedure. First, in-plane T1 images were registered to the mean functional image. Next, high-resolution T1 images were registered to the in-plane image. Structural images were then skull-stripped and normalized to the skull-stripped MNI template provided by FSL (Oxford Centre for Functional MRI of the Brain).

Additional pre-processing steps were implemented for the functional connectivity analyses. Data were bandpass filtered between 0.06 and 0.12 Hz, detrended, and standardized. Using the *nilearn* package in Python 2.7, we extracted regional timeseries from 5mm spherical regions defined from a whole-brain atlas (Power et al., 2011) and regressed out the average timeseries in each individual’s white matter and cerebrospinal fluid, as well as six head motion parameters. We also censored frames with framewise displacement (FD) > 0.5 mm and excluded participants with more than 40% of frames censored. The results were unchanged when censored frames were included in analyses.

### Putative cognitive systems

Our work employed a well-studied parcellation of the brain with 264 regions of interest that were each comprised of a 5 mm sphere (Power et al., 2011). From this whole-brain parcellation, we identified the regions that were specifically involved in response inhibition by conducting a reverse inference meta-analysis using the term “response inhibition” in the Neurosyth database (Yarkoni, Poldrack, Nichols, Van Essen, & Wager, 2011). Similarly, to identify regions involved in self-processing and mentalizing, we conducted two additional reverse inference meta-analyses using the terms “self-referential” and “mentalizing,” respectively. For each meta-analysis, we identified studies that matched the key phrase (threshold = 0.001, results of database query as of October 2017), yielding 176 studies related to response inhibition, 127 studies related to self-processing, and 124 studies related to mentalizing. For each set of studies, we submitted the associated MNI coordinates to a Neurosynth meta-analysis and saved the FDR-corrected (p<0.01) reverse inference map.

Regions of interest were considered to be involved in the process of interest if at least half of the voxels in the region were significantly activated in the FDR-corrected reverse inference map for that term. Interestingly, we found no overlap between regions involved in response inhibition and regions involved in self-processing or mentalizing as defined by the Neurosynth meta-analyses. We did, however, find overlap between regions involved in self-processing and mentalizing systems; to allow us to differentiate between these two terms, we excluded 12 regions involved in both processes. All analyses reported in this study include 21 mentalizing regions (which we refer to collectively as the mentalizing system), 8 self-processing regions (which we refer to collectively as the self-processing system), and 13 response inhibition regions (which we refer to collectively as the response inhibition system; see Figure 1 panel A).

### Statistical analysis of imaging data

In our analyses, we focused on three markers of neurophysiological dynamics. First, we measured the average BOLD activation of each system during correct No-Go versus correct Go trials, as a marker of neural processes related to successful inhibition of prepotent responses. Second, we measured the functional connectivity within each system, operationalized as the average Pearson correlation coefficient estimated between time series of any two nodes in the system. Third, we measured the functional connectivity between systems, operationalized as the average Pearson correlation coefficient estimated between time series of any node in one system and any node in another system. We tested whether any of these three markers was associated with the behavioral measure of response inhibition efficiency, as well as whether these markers of neurophysiological dynamics might interact with one another to promote more efficient response inhibition.

### System activation

Our first set of analyses examined mean activation for correct No-Go versus correct Go trials to investigate neural activity associated with successful response inhibition. Using a general linear model implemented in SPM8, the voxel activity was predicted from weighted beta coefficients for BOLD activity during correct No-Go trials, false-alarm No-Go trials, and missed Go trials. The correct Go trials were treated as an implicit baseline condition. Next, we computed mean activity for each of the three cognitive systems of theoretical interest: response inhibition, self-processing, and mentalizing systems. We extracted the contrast weight coefficients for the correct No-Go trials versus correct Go trials from each of the 42 regions of interest identified as a member of one of the three systems (Figure 1 panel A), and we averaged the response across each region in the system to calculate mean system activity.

We conducted three subsets of analyses using the mean activity of each system to test our planned hypotheses. The first subset investigated whether the system was recruited during successful inhibition of prepotent responses. Using one-sample *t*-tests, we assessed whether average activation in the response-inhibition system, self-processing system, and mentalizing system was stronger for correct No-Go versus correct Go trials. In the second subset of tests, we more directly assessed the system’s role in task performance variability. Using a separate model for each of the three systems, we computed the correlation between average system activity and the Go/No-Go efficiency score that accounts for the speed/accuracy trade-off during task performance (see Eq. 1). Finally, the third subset of tests employed multiple regression analyses to examine the interaction between systems and task performance. Given multicollinearity between the self-processing and mentalizing systems, we built two models using the Go/No-Go efficiency score as the dependent variable: (i) one model used mean activation in the response inhibition system and mean activation in the self-processing system as independent variables, whereas (ii) the other used mean activity in the response inhibition and mentalizing systems as independent variables.

### Functional Connectivity

Our second set of analyses examined similar questions as the mean activation analyses, but here, the cognitive system activity was characterized using functional connectivity to examine within-and between-system communication. Using the same 42 regions that comprise the three cognitive systems (Figure 1 panel A), we computed the Pearson correlation coefficient between the time series of each pair of regions separately for each run for each participant. We constructed a 42 × 42 functional connectivity matrix where the *ij*^th^ element of the matrix represented the correlation coefficient between the activity time series of region *i* and the activity time series of region *j*. We then averaged the functional connectivity matrices for the two runs for each participant to yield a single functional connectivity matrix for each participant. We then averaged the connectivity across all regions within each of the three cognitive systems to compute each system’s within-system connectivity. Finally, we computed two between-system connectivity values as the Pearson correlation coefficient between the response inhibition system and each of the two social brain systems (response inhibition x mentalizing, response inhibition x self-processing).

We then used both within-and between-system connectivity in two subsets of hypothesis-driven analyses. The first subset mirrored those executed using mean system activation and examined whether connectivity was associated with task performance variability. Using a separate model for each of the three systems, we computed the correlation between the Go/No-Go efficiency score and within-system connectivity. Two additional models assessed the correlation between the Go/No-Go efficiency score and each of the between-system connectivity values. The second and final subset of analyses directly tested whether functional connectivity might compensate for weaker mean activity in an adolescent’s response inhibition system to preserve successful task performance. To test this hypothesis, we ran 5 separate multiple regression models using the Go/No-Go efficiency score as the dependent variable: the first three had mean activation in the response inhibition system and one of the within-system connectivity as the independent variables, whereas the other two had mean activation in the response inhibition system and one of the between-system connectivity as the independent variables.

### Social Network Moderation Analyses

Finally, we investigated the relationship between an adolescent’s real-life social network structure and their neural and behavioral responses during a response inhibition task. Following similar logic as the mean activation and connectivity analyses, we employed each of the three properties of their real-life social network in two subsets of analyses. Separate models were employed throughout this set of comparisons based on the multicollinearity between activation across the three cognitive systems (i.e., individuals with greater activation in one system tended to have greater activation in other systems) as well as the multicollinearity among the social network metrics (e.g., individuals with a larger social network tended to have more communities within the social network).

The first subset of analyses examined the relationship between each network property and the neural response during task performance. A correlation was computed for each combination of network property metric (number, size, and modularity of communities) and brain measurement (mean activity of each system, within-connectivity of each system, and the two between-system connectivity values). The second and final subset of analyses assessed whether any of the three social network properties moderated the observed relationship between brain activity and task performance. Separate regression models were run with the adolescents’ Go/No-Go efficiency score as the dependent variable and each combination of brain measurement (mean activity of each system, within-connectivity of each system, and the two between-system connectivity values) and social network property (number, size, and modularity of communities) as the independent variables.

## Acknowledgements

S.H.T. and J.M.V. acknowledge mission funding from the U.S. Army Research Laboratory, including work under contract W911NF-11-2-0030. D.S.B. acknowledges support from the John D. and Catherine T. MacArthur Foundation, the Alfred P. Sloan Foundation, the ISI Foundation, the Paul Allen Foundation, the U.S. Army Research Laboratory (CaNCTA-W911NF-10-2-0022, Bassett-W911NF-14-1-0679, Grafton-W911NF-16-1-0474, and DCIST-W911NF-17-2-0181), the Office of Naval Research, the National Institute of Mental Health (2-R01-DC-009209-11, R01 – MH112847, R01-MH107235, R21-M MH-106799), the National Institute of Child Health and Human Development (1R01HD086888-01), National Institute of Neurological Disorders and Stroke (R01 NS099348), and the National Science Foundation (BCS-1441502, BCS-1430087, NSF PHY-1554488 and BCS-1631550). Data collection was supported by the intramural research program of the Eunice Kennedy Shriver National Institute of Child Health and Human Development contract #HHSN275201000007C (PI: Bingham); a University of Michigan Injury Center Pilot Grant (PI: Falk); an NIH Director’s New Innovator Award #1DP2DA03515601 (PI: Falk) and NIH/NICHD IR21HD073549-01A1 (PI:Falk). The authors gratefully acknowledge the University of Michigan Transportation Research Institute for research assistance; the staff of the University of Michigan fMRI Center; and Raymond Bingham, Jean Shope, Marie Claude Ouimet, Anuj Pradhan, Bruce Simons-Morton, Kristin Shumaker, Elizabeth Beard, Jennifer LaRose, Farideh Almani, Andrea I. Barretto, Alyssa Templar and Johanna Dolle. The content is solely the responsibility of the authors and does not necessarily represent the official views of any of the funding agencies.

